# Soybean drought resilience: contributions of a brassinosteroid functional analogue

**DOI:** 10.1101/742429

**Authors:** Lucia Sandra Pérez-Borroto, Laila Toum, Atilio Pedro Castagnaro, Justo Lorenzo Gonzalez-Olmedo, Francisco Coll-Manchado, Björn Gunnar Viking Welin, Yamilet Coll-García, Esteban Mariano Pardo

## Abstract

Drought is one of the most important causes of severe yield loss in soybean worldwide, threatening food production for the coming years. Phytohormones such as brassinosteroids can increase response to water deficit. However, natural brassinosteroids low stability precludes large-scale field application, challenging research and development of more stable and cost-effective analogues. Seeking functional analogues capable of improving plant drought-response, we investigated for the first time the effect of DI-31 in Arabidopsis and soybean. We found that, in *A. thaliana,* the DI-31 increased root growth, biomass accumulation, leaf number *per* plant, triggered antioxidant response and dose-dependent stomatal closure, requiring NADPH and peroxidase-dependent ROS production. In soybean, the relative water content, water use efficiency, biomass production and duration, root length, free proline, chlorophyll and carotenoid accumulation and enzymatic antioxidants activity were stimulated by DI-31 application after four and eight days of mild water shortage, while significantly reduced the lipid-peroxides content. Additionally, our results demonstrated that DI-31 diminishes the nodular senescence and successfully maintains the N homeostasis through a fine tune of biological/assimilative N2-fixation pathways. These findings support the DI-31 potential use as a sustainable alternative for integrative soybean resilience management under drought.

**Highlight:** Brassinosteroid analogue DI-31 improves soybean growth, water economy, respiration, anti-stress response and nitrogen homeostasis under drought. Thus, they may be considered as a sustainable and environmentally-safe alternative for raising legumes climate resilience.

## Introduction

Brassinosteroids (BRs) are a group of growth-promoting plant hormones, isolated and characterised from canola pollen (*Brassica napus* L.) (Moreno-Castillo et al. 2018). Up to date, nearly 70 kinds of natural BRs analogues have been isolated from tissues of various plant species (Tang et al. 2016). BRs possess beneficial pleiotropic effects due to its broad and highly coordinated cell modulation capacity (Sahni et al. 2016). They influence various developmental processes and agronomical traits (Gonzalez-Olmedo et al. 2004; González-Olmedo et al. 2005; Vriet et al. 2012). Moreover, BRs can increase plant response to a wide range of stresses (Moreno-Castillo et al. 2018). That is why researchers have been exploring strategies directed to increase crops yields and stress tolerance/resistance, like BRs exogenous application and the genetic manipulation of its endogenous levels. However, the BRs low stability in the field precludes large-scale application (Sakai et al. 1999), being replaced by functional or structural analogues with higher biological activity and field average life (Sasse 2003). The use of BRs analogues with a predominantly growth-promoting effect in a wide range of plant species constitutes an alternative directed to improving crop yields (Rao et al. 2002). The most commonly used is 24-epibrassinolide despite its high production fees (Moreno-Castillo et al. 2018), so the study of other analogue molecules with higher activity and lower costs constitute a promising alternative.

Among abiotic stresses, water deficit has the most severe effect in worldwide agriculture and shows detrimental effects on plants stomatal morphology, photosynthesis, growth rate and oxidation-reduction balance (Chai et al. 2016). Soybean (*Glycine ma*x (L.) Merrill) is one of the socio-economic crops affected by drought. Considered the most worldwide cropped legume, soybean is an essential source of oil and protein (Hungria and Mendes 2015; Wang et al. 2018). However, the occurrence of water scarcity periods significantly reduced the crop photosynthesis, growth and nitrogen (N) fixation, causing grain quality and yield losses (Jin et al. 2018). To mitigate the detrimental effects of water deprivation, researchers develop strategies such as genotypes selection, identification of tolerant genetic sources and establishment of cultural management practices, like bio-stimulant applications.

The synthetic spirostanic BRs functional analogue (25R)-3β,5α-dihydroxy-spirostan-6-one (DI-31) property of the CEPN (Center of Studies of Natural Products, Chemistry Faculty, Havana University) is characterized by the presence of a spiroketalic ring instead of the typical BR side chain (Coll et al. 1995; Mazorra et al. 2002) and BR-like activity (Furio et al. 2018). Up to date, the DI-31 has been studied only as a component of the commercial bio-stimulant Biobras-16, with beneficial effects on photosynthetic rate and yield of greenhouse-grown pepper (*Capsicum annuum* L.) (Serna et al. 2012)and endive (*Cichorium endivia* L.) plants (Serna et al. 2013). Similarly, Biobras-16 application prevented the negative effect of salt stress in rice and lettuce plants (Serna et al. 2015).

Structurally, the DI-31 has an epoxy-oximic polar group, one of its major conformations, that interacts with the BRI1 BRs plant receptor with higher affinity and lower binding energy than 24-epibrassinolide (Moreno-Castillo et al. 2018), so it has a higher potential activity. Therefore, it could be useful to incorporate the DI-31 in a crop management strategy focused on increasing crop yield and to diminish the effect of abiotic stresses like osmotic imbalance and particularly drought. Besides, the application of hormonal molecules such as DI-31 could constitute an eco-friendly complement that stimulates soybean growth and N fixation maintenance under water scarcity. Nevertheless, in order to characterize this compound, it is very convenient to test its effects in a model plant such as *Arabidopsis thaliana* (L.) Heynh, and then to evaluate its potential in soybean plants.

In this study, we investigated for the first time the DI-31 molecule effect on *A. thaliana* growth, stomatal movement, oxidative burst and antioxidant enzymatic activity. Subsequently, we characterised the DI-31 action in soybean plants under water deficit for potential drought resilience, evaluating the compound effect on photosynthesis, water economy, biomass production and particularly N homeostasis under drought. These results support the potential use of this compound to crop management under drought conditions.

## Material and methods

### Plant material and growth conditions

All the experiments were conducted at the Estación Experimental Agroindustrial Obispo Colombres (EEAOC), Las Talitas, Tucumán, Argentina (S26°50’, W65°12’). The assays in wild type (WT) *A. thaliana* were developed using Columbia (Col-0) ecotype seeds. Seeds were disinfected for 5 minutes in a mixture of commercial bleach, distilled water and ethanol (1:1:8). Subsequently, they were washed three times with 96% ethanol under sterile conditions and seeded in Petri dishes containing MS medium (Murashige and Skoog 1962), with 1% (w/v) sucrose, 0.5 g/L of MES pH 5.8 (Duchefa, Holland) and 0.8% (w/v) agar (Sigma, USA). Seeds in the plates were stratified in the dark for three days at 4°C and then transferred to a growth chamber at 22-23°C temperature and 16 h light-8 h darkness photoperiod. Seed germination was monitored for five days, and then seedlings were transferred to new Petri dishes with fresh MS medium or plastic glasses (diameter: 5.5 cm, height: 9 cm) filled with GrowMix® Multipro commercial substrate (Terrafertil S.A., Argentina), according to each test requirements. Only the DI-31 curve dose-response assay was carried out in Petri dishes using 5-day-old seedlings transferred to plates with fresh medium and supplemented with the compound at different doses. The rest of the experiments were performed using 3-week-old plants (except stomatal assays, performed with 4-week-old plants) grown in plastic glasses filled with commercial substrate and irrigated with Hoagland Complete Solution (Hoagland and Arnon 1950).

The experiments in soybean were performed in glasshouse conditions using the commercial cv. Munasqa RR, selected from previous comparative trials with soybean cv. TJ2049 and MG/BR 46 Conquista (Perez-Borroto unpublished, ongoing). An amount of 1000 Munasqa RR homogeneous seeds (EEAOC Germplasm Bank), manually harvested and with a high germination potential were selected and planted in 4 L plastic pots (diameter: 18 cm, height: 21 cm) filled with GrowMix® Multipro commercial substrate. Before sowing, seed were inoculated with *Bradyrhizobium japonicum* E109 (9 × 10^9^ viable cells kg^−1^ of seeds) in order to guarantee maximum soybean plant performance. Four seeds *per* pot were placed to ensure germination. When plants reached V1 vegetative stage (open leaf at the unifoliate node) according to (Fehr et al. 1971), the number of plants *per* pot was reduced by half. Trials were performed using V3 (second open trefoil) or V5 (fourth open trefoil) vegetative stage plants. During the experiments, the environmental (ET) and substrate temperatures (ST), relative humidity (RH) and the photosynthetically active radiation (PAR) were measured every 15 min, averaged and recorded at one h intervals with data loggers (Cavadevices.com, Buenos Aires, Argentina). The environmental variables evaluated in the period of experiments presented the following average values: ET = 28°C (± 7°C); ST = 22.4°C (± 4°C); RH = 90.2%; PAR = 648.37 μmol m^-2^s^-1^. Plants grew with a 12 h average photoperiod. Pots were distributed in a completely randomized design and weekly moved and rearranged to minimize possible environmental effects.

### Water availability treatments

The estimation of substrate water content (SWC) in each pot was performed as described by (Pereyra-Irujo et al. 2012). Briefly, the weight of each empty pot and the commercial substrate at the beginning of the experiment were determined. Also, we quantified the fresh weight of the plants every five days. Then, the data was used to calculate the water content of each pot and the amount of water that had to be added every day to reach the desired SWC. Subsequently, the relationship between SWC and substrate water potential (Ψs) was determined according to (Richards 1965). All pots were watered to a 22% SWC corresponding to a Ψs of -0.05 MPa until the imposition of water deficit treatments. Stress imposition was performed according to (Pardo et al. 2015) drought phenotyping protocol, which consists in maintaining the SWC at 14% corresponding to a Ψs of -0.65 MPa during eight days. The Ψs corresponding to the water deficit treatment was reached in a 1-2 days interval. The plant’s water status (relative water content) was monitored throughout the water shortage period to ensure stress occurrence. Additionally, the experimental design also included six pots filled with substrate distributed in the two SWC treatments (Ψs of -0.05 and -0.65 MPa). The pots were distributed randomly and daily watered and weighed to quantify the amount of water evaporated from the substrate.

### DI-31 dose-response curve and growth promotion

5-day-old WT *A. thaliana* seedlings were transferred to new Petri dishes and divided in four groups (10 plants *per* treatment): (i) MS medium (control treatment), (ii) MS medium + DI-31 (0.22 µM; 0.1 mg/L), (iii) MS medium + DI-31 (1.12 µM; 0.5 mg/L) and (iv) MS medium + DI-31 (2.23 µM; 1 mg/L). Five days after treatments, we photographed the seedlings and measured morphological growth indicators such as the number of leaves, root length and biomass production. The DI-31 concentration of 2.23 µM was chosen to perform further assays. The experiment was performed twice with similar results.

### Stomatal measurements

Stomatal assays (Gudesblat et al. 2009) were carried out using *A. thaliana* 4-week-old plants (non-flowered). We placed the epidermal peels in well plates with 500 µL of a 10:10 buffer solution (10 mM KCl and 10 mM MES-KOH, pH = 6.15) under the normal culture conditions for 2 h. Then, different treatments were applied and the epidermis were incubated for an additional 1.5 h. To assess the DI-31 effect on stomata four treatments were defined: (i) untreated buffer solution, (ii) buffer solution + ABA (20 µM), (iii) serial dilutions of 24-epibrassinolide and (iv) DI-31 (0.1; 0.5; 1; 5; 10 and 20 µM). To test whether DI-31-induced stomatal closure is dependent on ROS production, specific inhibitory compounds treatments were defined: (i) Diphenyleneiodonium (DPI) 10 μM (Sigma, USA) + ABA (20 µM), (ii) DPI + DI-31 (10 µM), (iii) Salicylichydroxamic acid (SHAM) 2mM (Sigma, USA) + ABA (20 µM) and (iv) SHAM acid + DI-31 (10 µM). DPI and SHAM were incubated 30 min before DI-31 treatments. We performed 40 measurements *per* treatment and presented the data as the average of 80 measurements collected from four independent experiments.

### Oxidative burst assay

To determine whether DI-31 can activate defence mechanisms as the respiratory burst in well-watered *A. thaliana* plants, we evaluated the appearance of superoxide radicals. Three-week-old WT plants were grouped according to the following treatments: (i) control with distilled water (DW) and (ii) DI-31 (2.23 µM). Treatments were foliarly applied (by sprinkling) to the plant rosettes until they reached the drip point. To determine the oxidative burst, we collected three plants for each treatment and harvest timing (total of 24 plants). From each plant collected, the sixth and seventh rosette leaves were detached at 6, 12, 24 and 48 h after DI-31 treatment and subjected to NBT (Nitroblue tetrazolium) staining protocol (Doke 1983). We selected the sixth and seventh leaves (counting from the youngest leaf) representing fully expanded leaves.

### Antioxidant response measurements

WT *A. thaliana* plants were used to assess the DI-31 ability to stimulate enzymatic antioxidants activity such as the superoxide dismutase (SOD, EC 1.15.1.1) (Li 2012), catalase (CAT, EC 1.11.1.6) (Chance and Maehly 1955), ascorbate (APX, EC 1.11.1.11) (Nakano and Asada 1987) and phenol peroxidase (POX, EC 1.11.1.7) (Kar and Mishra 1976) as well as the protein accumulation (Bradford 1976). We performed an enzymatic uniform extraction (Liu et al. 2010; Singh et al. 2014). The DI-31 (2.23 µM) was sprinkled to drip point. Two experiments were performed in a growth chamber using 3-week-old plants, distributed in two treatments: (i) well-irrigated plants sprinkled with distilled water (control) and (ii) well-irrigated plants sprinkled with DI-31. Ten plants *per* treatment were collected and sampling times (60 plants *per* experiment) were at 0, 24 and 48 hours after the compound application.

Subsequently, we determined the effect of DI-31 foliar application in soybean antioxidant activity under well-irrigated and water scarcity conditions in two glasshouse experiments. When the Munasqa RR plants reached the V3 phenological stage were distributed into four groups corresponding to the following treatments: (i) well-irrigated plants (Ψs= −0.05 MPa) sprinkled with distilled water (DW), (ii) well-irrigated plants sprinkled with DI-31, (iii) stressed plants (Ψs= −0.65 MPa) sprinkled with DW and (iv) stressed plants sprinkled with DI-31. Then, we induced the water deficit in the treatments (iii) and (iv). Once the treatments reached the SWC and Ψs corresponding to moderate water stress, the DW and the DI-31 (2.23 µM) were applied using the same method previously described for *A. thaliana* trials. Ten soybean plants *per* treatment were used and V2, V3 and V4 leaves from each plant (total of 120 plants *per* experiment) were collected at 0, 4 and 8 days after the DW and DI-31 application. Besides the antioxidant enzymes and protein measurements, the content of MDA (Hodges et al. 1999), free proline (Bates et al. 1973), chlorophyll (Porra 2002) and carotenoids (Riemann 1978) were determined.

### Soybean growth, water economy and N fixation measurements

The experiments were carried out with V5 Munasqa RR plants (20 plants *per* treatment and sampling time, a total of 240 plants *per* experiment), distributed into the same four groups corresponding to the previously described treatments. We performed the stress imposition and DW and DI-31 application, as we described in the previous section. At 0, 4 and 8 days after the application of the compound, the whole plants were collected. For each sampling time, 15 collected plants were used to determine morphophysiological characters associated with growth, nodulation and N fixation. The remaining five plants, collected at each time, were used to determine the water status according to the (Weatherley 1950) relative water content (RWC) method. The water use efficiency (WUE), defined as the ratio between the above-ground biomass and the water consumed (Van Halsema and Vincent 2012) was also measured. As growth indicators, we quantified the biomass production (Porcel et al. 2003) and then calculated the biomass duration (BMD) over time (Hunt 1978). The number of leaves, stem length and thickness and primary root length, were also quantified. As nodulation parameters, we collect the nodules located in the root crown (an imaginary cylinder of 2.5 cm of diameter and length) according to (Burton 1976). Subsequently, the nodules were cut to visualize their activity status according to the Leghemoglobin colouration, then were labelled and photographed together with a scale for further morphological analysis. The number of active nodules *per* plant was quantified. In order to determine if the DI-31 have any effect on nodules development under stressful conditions, morphological parameters such as equatorial and polar diameter, the thickness of periderm and cortex area (outer, middle and inner) and the estimated area of the infected central medulla were measured in all the active nodules (Kanu and Dakora 2017), using the image processing and analysis program *ImageJ* (version 1.52). The N fixation parameters were measured in extracts obtained from the aerial portion of the plants. Indicators such as the *in vivo* activity of Nitrate Reductase (NR) enzyme (Jaworski 1971), nitrate (Cataldo et al. 1974), α-amino acids (Herridge et al. 1990) and ureide content (Young and Conway 1942) were determined. Additionally, we calculated the ureide relative abundance (Takahashi et al. 1992) and the percentage of biological N fixed (Herridge et al. 1990). The experiments were performed twice with similar results.

### Statistical analysis

All data were analyzed in GraphPad Prism 5.01, using ANOVA and Tukey’s test (Tukey’s HSD). Each treatment value is presented as the arithmetic mean ± S.E. (standard error) marked with letters in the graphs.

## Results

### DI-31 enhances A. thaliana growth and triggers antioxidant response

To assess DI-31 effect on growth, we measured root length, the number of leaves and biomass increase in *A. thaliana* WT plants (Fig. 1). The results showed that DI-31 promotes root length in a dose-dependent manner, reaching a ∼46% length increase in the plants treated with the highest concentration (Fig. 1**b**). Furthermore, compared to the control and the DI-31 lowest dose (0.22 µM), the treatments with DI-31 highest concentrations (1.12 and 2.23 µM) showed a significant effect on the number of leaves *per* plant (Fig. 1**c**) and biomass accumulation (Fig. 1**d**), five days after the compound application.

**Fig. 1.**
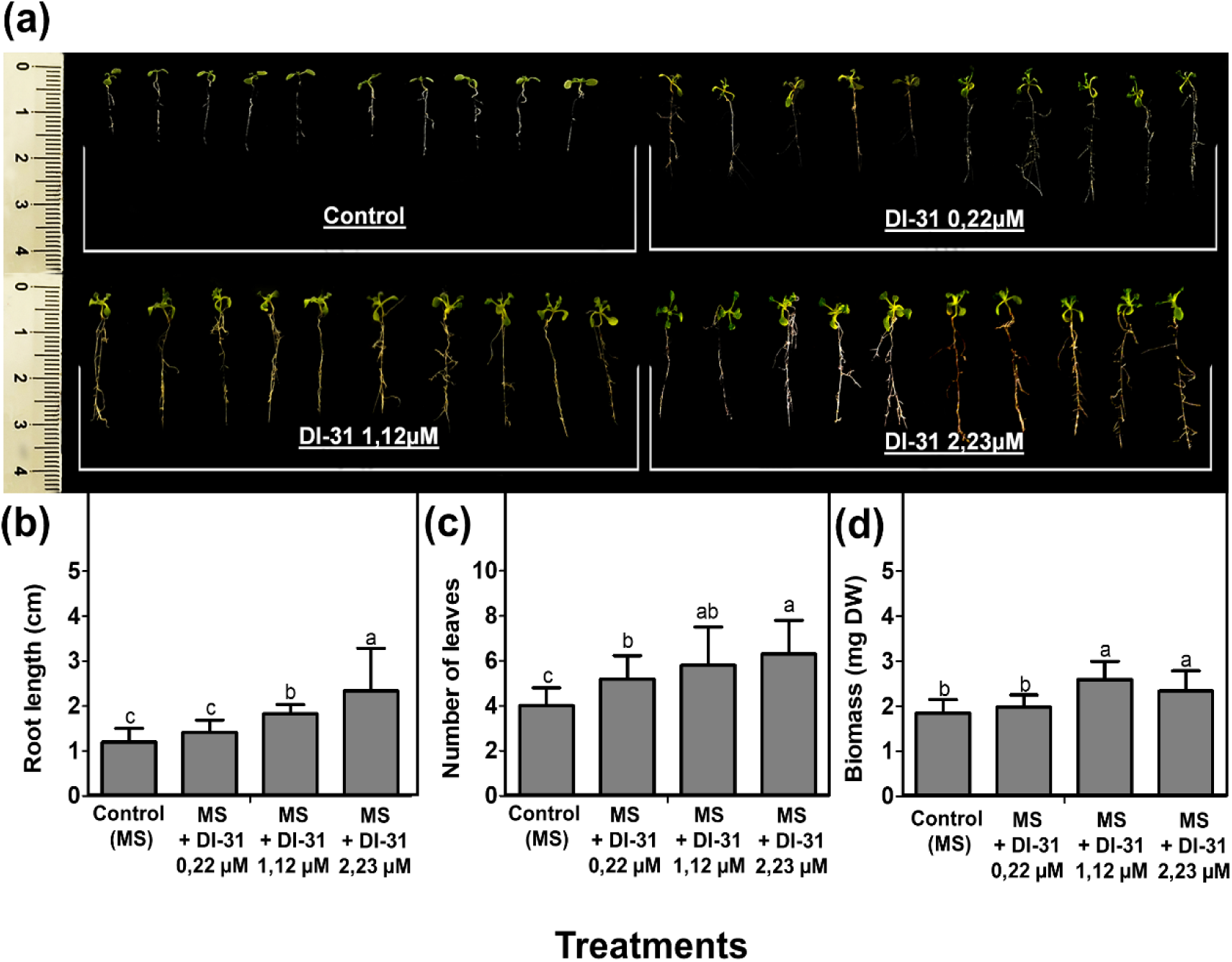
DI-31 application promotes *A. thaliana* Col-0 growth. (**a**) Representative image of plants growing proportionally to DI-31 concentration (0.22 µM (0.1 mg/L), 1.12 µM (0.5 mg/L) and 2.23 µM (1 mg/L)). Morphological parameters such as (**b**) root length (cm), (**c**) number of leaves and (**d**) biomass increase (mg) measured in 10-day-old plants grown in MS medium with and without DI-31 at different doses. Data represent the mean (±SD) of two independent experiments (n=80). Different letters on top of the bars indicate significant difference as determined by ANOVA with *post hoc* contrasts by Tukey’s test: (P<0.05).

Furthermore, the DI-31 effect on enzymatic antioxidants activity over time was assessed and compared to control. The DI-31 application progressively stimulated the activity of the SOD (Fig. 2**a**), APX (Fig. 2**b**) and POX (Fig. 2**c**) enzymes. The POX reached the highest activity value 24 h after the compound application, while the SOD and APX did it at 48 h. CAT enzyme remains unmodified (Fig. 2**d**). The protein content (Fig. 2**e**) did not statistically differ among the treatments and harvest timings. Additionally, we assessed the DI-31 effect on superoxide radicals production (Fig. 2**f**). After 6 hours, was detected the appearance of respiratory burst symptoms in DI-31-treated rosettes, which reached the highest formation of blue formazan points 48 h after the compound application.

**Fig. 2.**
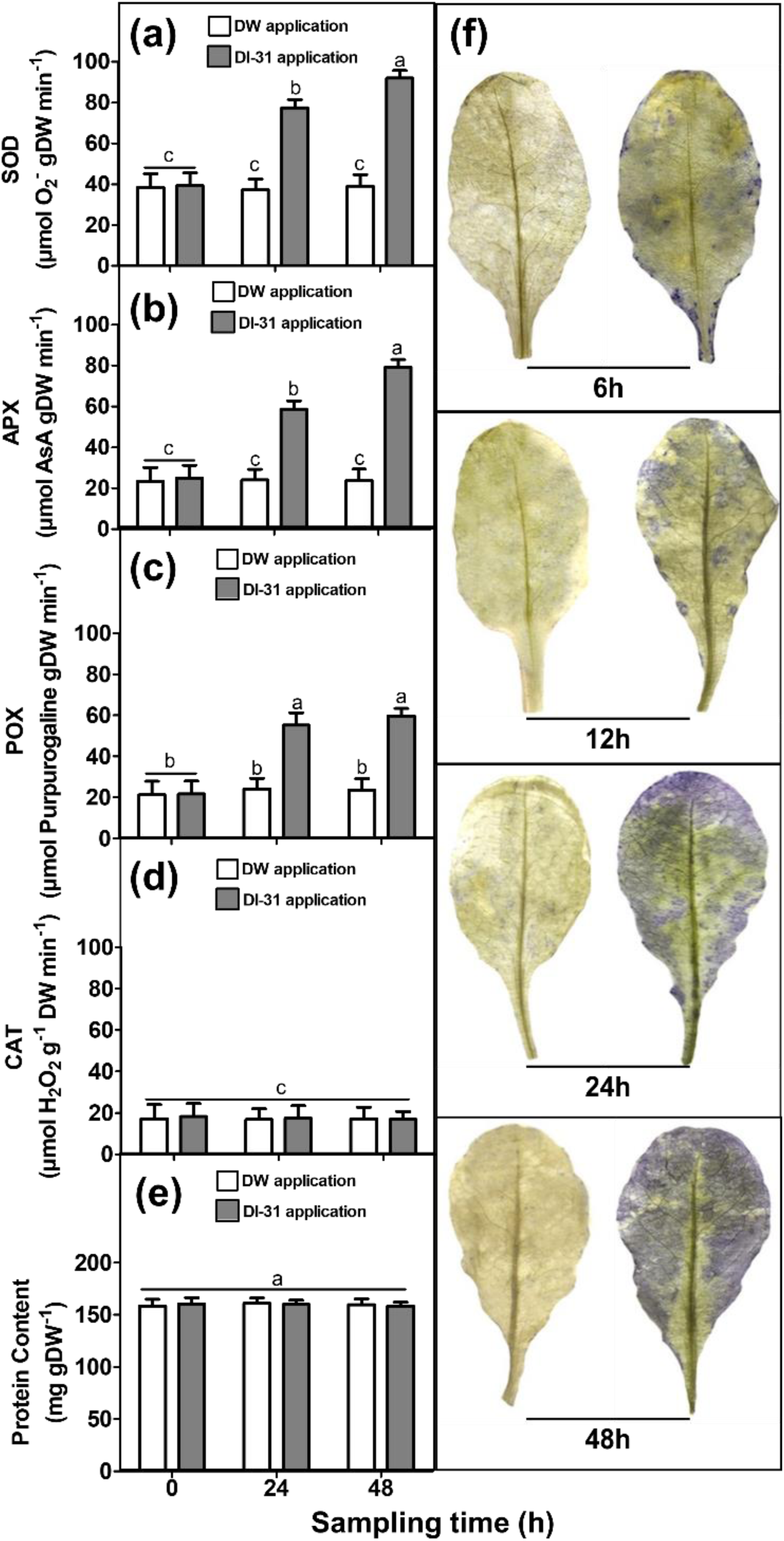
DI-31 application stimulates the respiratory burst and antioxidant response of *A. thaliana* Columbia 0 plants under well-watered conditions. (**a**) Superoxide dismutase (SOD), (**b**) Ascorbate peroxidase (APX), (**c**) Phenol peroxidase (POX) and (**d**) Catalase (CAT) enzymatic activities, and (**e**) protein content determined in WT plants leaves (n=120) collected at the trial beginning (0), 24 and 48 h after distilled water (DW) and DI-31 (2.23 µM) foliar applications. (**f)** Representative image of the oxidative burst induced by DI-31 application, measured through superoxide radicals accumulation by NBT staining on the sixth and seventh plant leaves, collected at 6, 12, 24 and 48 h after treatment (n=24). Different letters on top of the bars indicate significant difference as determined by ANOVA with *post hoc* contrasts by Tukey’s test: (P<0.05).

### DI-31-mediated stomatal closure requires ROS production in A. thaliana

As it was previously described for 24-epibrassinolide (Shi et al. 2015), we decided to test whether DI-31 can induce stomatal closure. Stomatal closure patterns induced by 24-epibrassinolide and DI-31 were very similar in all treatments applied. DI-31 significantly induced stomatal closure in a dose-dependent manner similarly to 24-epibrassinolide (Fig. 3**a**), in agreement with previous results.

**Fig. 3.**
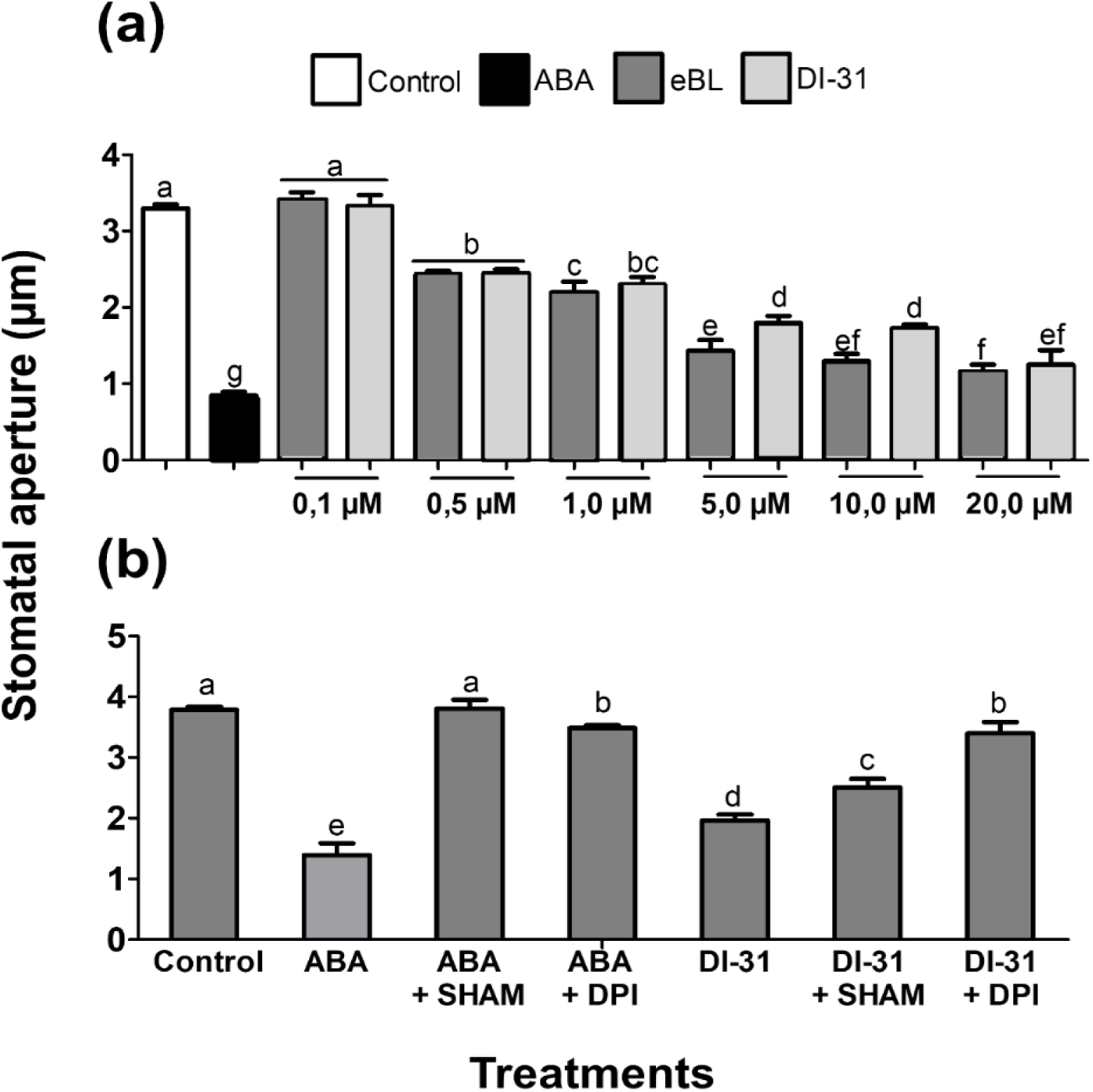
DI-31 induce *A. thaliana* dose-dependent stomatal closure, requiring NADPH and peroxidase-dependent ROS production. (**a**) DI-31 and 24-epibrassinolide stomatal closure comparative experiments in 4-week-old plants, using an opening control (untreated KCl-MES-KOH buffer), a closing control with ABA (20 μM) and treatments with serial dilutions (0.1; 0.5; 1; 5; 10 and 20 μM) of DI-31 and 24-epibrassinolide. Stomatal apertures were measured 1.5 hours after the compound application. Data represent the mean (± SE) of two independent experiments (40 stomata *per* treatment, a total of 560 stomata). (**b**) Stomatal closure assay with DPI and SHAM ABA inhibitors carried out in 4-week-old plants, using an opening control (untreated KCl-MES-KOH buffer), a closing control with ABA (20 μM) and treatments with specific inhibitors of NADPH-oxidases (DPI: 10 μM) and cell-wall peroxidases (SHAM: 2 mM) with and without DI-31 (10 µM). Stomatal apertures were measured 1.5 hours after the application of the treatments. Data represent the mean (± SE) of two independent experiments (n=480 stomata). Different letters on top of the bars indicate significant difference as determined by ANOVA with *post hoc* contrasts by Tukey’s test: (P<0.05).

We additionally conducted a stomatal assay to test whether inhibition of ROS could impair the DI-31-mediated stomatal closure. For this purpose, we used DPI to inhibit NADPH oxidases and SHAM for suppressing cell-wall peroxidase activity. Our results showed that DI-31 stomatal closure was partially inhibited by both compounds, being more remarkable the effect of DPI. These results suggest that inhibition of NADPH oxidases prevents DI-31 from promoting stomatal closure and, to a lesser extent, peroxidases are also required (Fig 3**b**) for DI-31 action on stomata.

### DI-31 enhances soybean growth, water economy and stress response under drought

To validate DI-31 effect observed in *A. thaliana*, we decided to test this compound in a crop of agronomic importance. Thus, we conducted experiments in soybean cv. Munasqa RR. A wide range of morphophysiological and biochemical parameters associated with growth, water economy and stress response under water shortage were measured. Our results showed the protective effect of DI-31 in soybean plants subjected to water scarcity. The stressed plants treated with the compound showed an attenuated defoliation phenotype (Fig. 4**f**), evidencing still green and hydrated young leaves, with less typical drying symptoms such as curling. In agreement with these findings, our results indicated that application of DI-31 attenuated the RCW (Fig. 4**a**) and WUE (Fig. 4**b**) reduction under stress. Well-watered plants (both with DW and DI-31 application) evidenced a RWC of ∼91% during the experiment. Otherwise, under water shortage, DI-31-treated plants exhibited RWC values of 83 and 76.2%, while untreated plants showed 70.5 and 58.3%, on the fourth and eighth day of the trial, respectively.

**Fig. 4.**
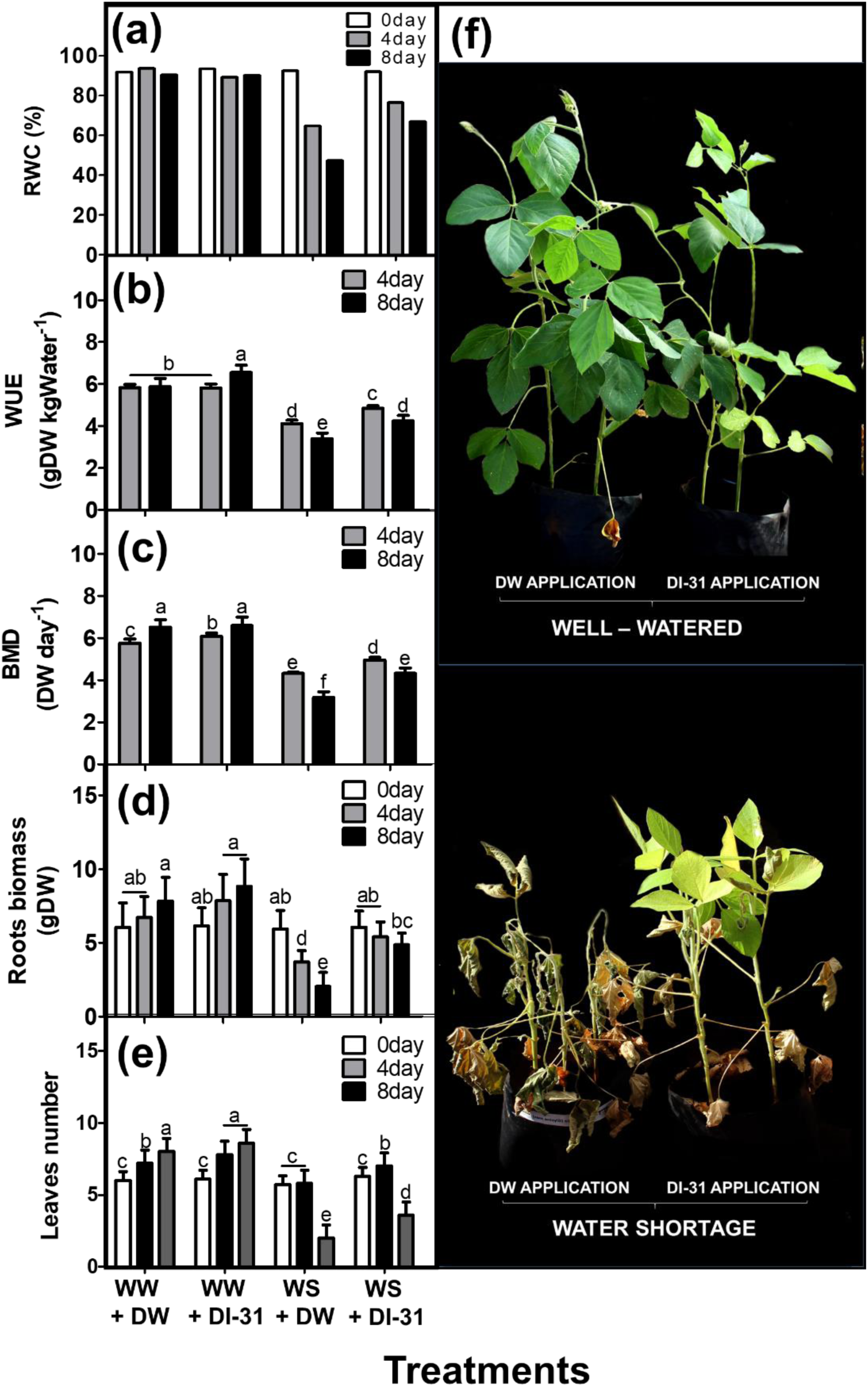
DI-31 contributes to soybean growth and water preservation under water shortage. Morphophysiological parameters such as (**a**) plant relative water content (RCW), (**b**) water use efficiency, (**c**) biomass duration, (**d**) roots biomass and (**e**) leaves number, were measured in V5 soybean cv. Munasqa RR plants. (**f**) Representative image of the highly contrasting phenotype observed in the Munasqa RR plants treated with distilled water (DW) and DI-31 and submitted to eight days of water shortage. Four treatments were defined: (i) well-irrigated plants (substrate water potential (Ψs) = −0.05 MPa) sprinkled with DW, (ii) well-irrigated plants sprinkled with DI-31, (iii) stressed plants (Ψs= −0.65 MPa) sprinkled with DW and (iv) stressed plants sprinkled with DI-31. Once the stress treatments reached the substrate water content and Ψs corresponding to moderate water stress, the DI-31 (2.23 µM) and DW treatments were performed by sprinkling to the drip point. Whole plants were collected at 0, 4 and 8 days after the water shortage. Data represent the mean (± SE) of two independent experiments n= 60 (**a**-**b**) and n= 180 (**c-e**). Different letters on top of the bars indicate significant difference as determined by ANOVA with *post hoc* contrasts by Tukey’s test: (P<0.05).

On the other hand, at the fourth and eighth day of the experiment, the WUE, under well-watered conditions, increased 4.3 and 11.5% in DW and DI-31-treated plants, respectively. Under water shortage, plants with DW application showed a 26.4 and 42% of WUE reduction, in comparison with DW-treated and well-watered plants at the fourth and eighth day of the experiment. Meanwhile, during the same period, the stressed plants treated with DI-31 showed a 13.6 and 27% of WUE reduction, compared to the control plants. The BMD under drought was also favoured by DI-31, compared to DW-treated plants subjected to water scarcity (Fig. 4**c**). Other indicators associated with growth, such as stem thickness and length, remained unmodified. Meanwhile, the DI-31 treatment stimulated the biomass production in the root system (Fig. 4**d**) and leaves development (Fig. 4**e**) in well-irrigated plants during the first four days after application.

The compound application over time also stimulated the specific activity of SOD (Fig. 5**a**) and APX enzymes (Fig. 5**b**), while the POX (Fig. 5**c**) and CAT enzymes (Fig. 5**d**) remained unmodified. No variations in protein content were observed (Fig. 5**e**). Findings that agree with the results showed in *A. thaliana* plants. Under water deprivation, treatment with DI-31 partially prevented chlorophyll pigments degradation (Fig. 5f-h), increased the total carotenoids (Fig. 5**i**) and free proline content (Fig. 5**j**) and limited the MDA accumulation (Fig. 5**k**) in leaf tissue.

**Fig. 5.**
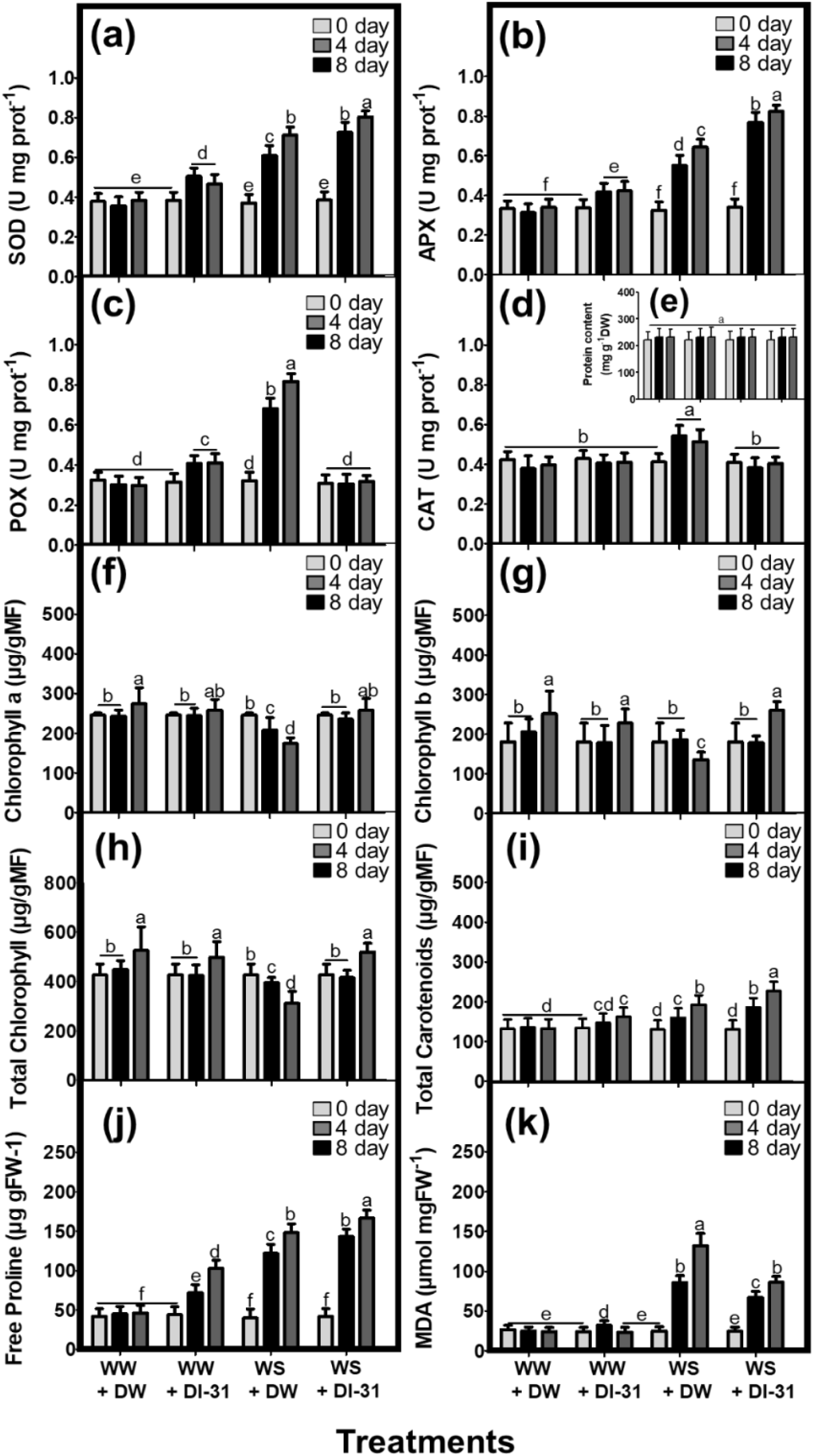
DI-31 application stimulates soybean anti-stress response under water shortage. Physiological and biochemical indicators such as (**a**) Superoxide dismutase (SOD), (**b**) Ascorbate peroxidase (APX), (**c**) Phenol peroxidase (POX) and (**d**) Catalase (CAT) specific activities, as well as the content of (**e**) protein, (**f**) chlorophyll a, (**g**) b and (**h**) total, (**i**) total carotenoids, (**j**) free proline and (**k**) malondialdehyde (MDA), where determined in V5 soybean cv. Munasqa RR plants. Four treatments were defined: (i) well-irrigated plants (substrate water potential (Ψs) = −0.05 MPa) sprinkled with distilled water (DW), (ii) well-irrigated plants sprinkled with DI-31, (iii) stressed plants (Ψs= −0.65 MPa) sprinkled with DW and (iv) stressed plants sprinkled with DI-31. Once the stress treatments reached the soil water content and water potential corresponding to moderate water stress, the DI-31 (2.23 µM) and DW treatments were performed by sprinkling to the drip point. The V2, V3 and V4 leaves from each plant were collected at 0, 4 and 8 days after the application of the treatments. Data represent the mean (± SE) of two independent experiments n= 240. Different letters on top of the bars indicate significant difference as determined by ANOVA with post hoc contrasts by Tukey’s test: (P<0.05).

### Nodulation and nitrogen homeostasis under drought and DI-31 treatments

Under water shortage, the treatment with DI-31 showed beneficial effects in Munasqa RR plants active/functional nodules (Fig. 6**b**) located in the imaginary root cylinder (Fig. 6**a**). During the first four days after the compound was applied, no significant decrease in the number of active nodules was observed, while at the eighth day we quantified a reduction of ∼31% (Fig. 6**d**). In contrast, DW-treated plants subjected to water shortage showed reductions in the number of active nodules by ∼53 and 57% on the fourth and eighth day of stress, respectively. From a more detailed analysis of the active nodules collected on the eighth day after DI-31 treatment (trial last day), significant differences were quantified in several morphological parameters (Fig. 6**c**). In well-watered and stressed plants, the compound application stimulated the infected central medulla estimated area, the equatorial and polar diameter of active nodules (Fig. 6e-g). While, in plants under water scarcity, significant reductions of these parameters were quantified, resulting in visually smaller nodules. Interestingly, under water shortage, the compound caused a thickening of the periderm (Fig. 6**h**) and a thinning of the nodular cortex area (Fig. 6**i**).

**Fig. 6.**
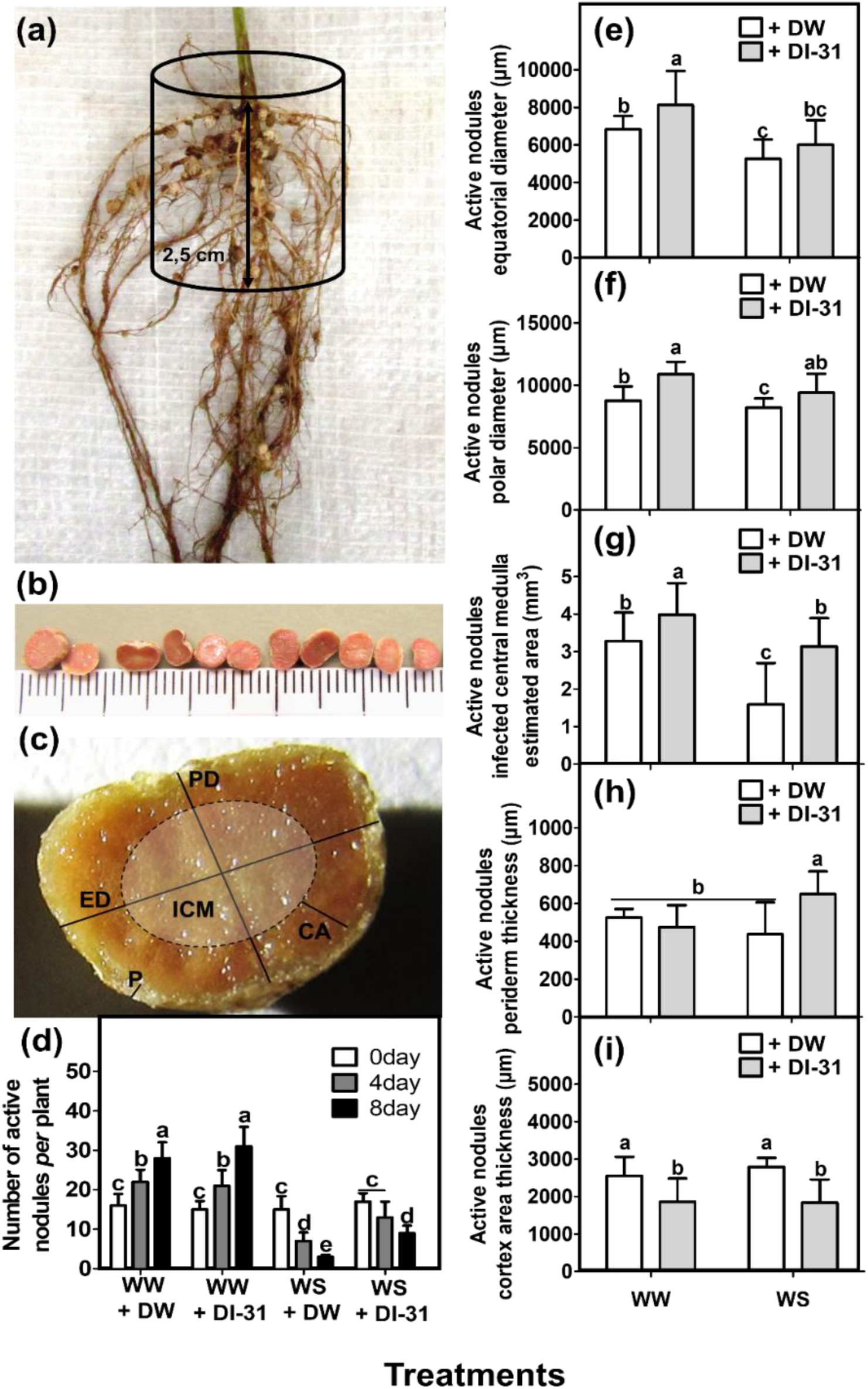
DI-31 treatment modulates soybean nodulation parameters under water shortage. Nodules located in the root crown imaginary cylinder (**a**) were collected and classified in active nodules according to the Leghemoglobin pink colouration (**b**), then the total of active nodules was quantified (**d**) and cut for morphological analysis (**c**). Nodulation parameters such as equatorial (**e**) and polar (**f**) diameter, the estimated area of the infected central medulla (**g**) and the thickness of periderm (**h**) and (**i**) cortex area (outer, middle and inner) were measured in V5 soybean cv. Munasqa RR active nodules using through image processing and analysis (*ImageJ* 1.52v). Four treatments were defined: (i) well-irrigated plants (substrate water potential (Ψs) = −0.05 MPa) sprinkled with distilled water (DW), (ii) well-irrigated plants sprinkled with DI-31, (iii) stressed plants (Ψs= −0.65 MPa) sprinkled with DW and (iv) stressed plants sprinkled with DI-31. Once the stress treatments reached the substrate water content and Ψs corresponding to moderate water stress, the DI-31 (2.23 µM) and DW treatments were performed by sprinkling to the drip point. Whole plants were collected at 0, 4 and 8 days after the application of the treatments. Data represent the mean (± SE) of two independent experiments (n= 180). Different letters on top of the bars indicate significant difference as determined by ANOVA with post hoc contrasts by Tukey’s test: (P<0.05).

The evaluated physiological and biochemical parameters associated with the N fixation showed significant alterations in the plants treated with the DI-31. Under well-watered conditions, the compound did not affect the *in vivo* NR activity (Fig. 7**a**) or the nitrate content (Fig. 7**b**); while the content of α-amino acids (Fig. 7**c**) increased at the fourth and eighth day after treatment. However, at the fourth day of water shortage, the *in vivo* NR activity and nitrate content increased due to DI-31 action, as well as the α-amino acids levels, that also increased at eighth days after treatment. Plants subjected to DW and water shortage treatment showed a constant increase of these three indicators. On the other hand, the ureide content in Munasqa RR leaves (Fig. 7**d**) significantly increased over time, in well-watered and stressed plants treated with DI-31. Finally, the compound application increased in ∼15% the ureides relative abundance (Fig. 7**e**) and ∼16% the percentage of biological N fixed (Fig. 7**f**) in well-watered plants, while Munasqa RR plants, submitted to water shortage and DI-31 treatments, showed the maintenance of both indicators.

**Fig. 7.**
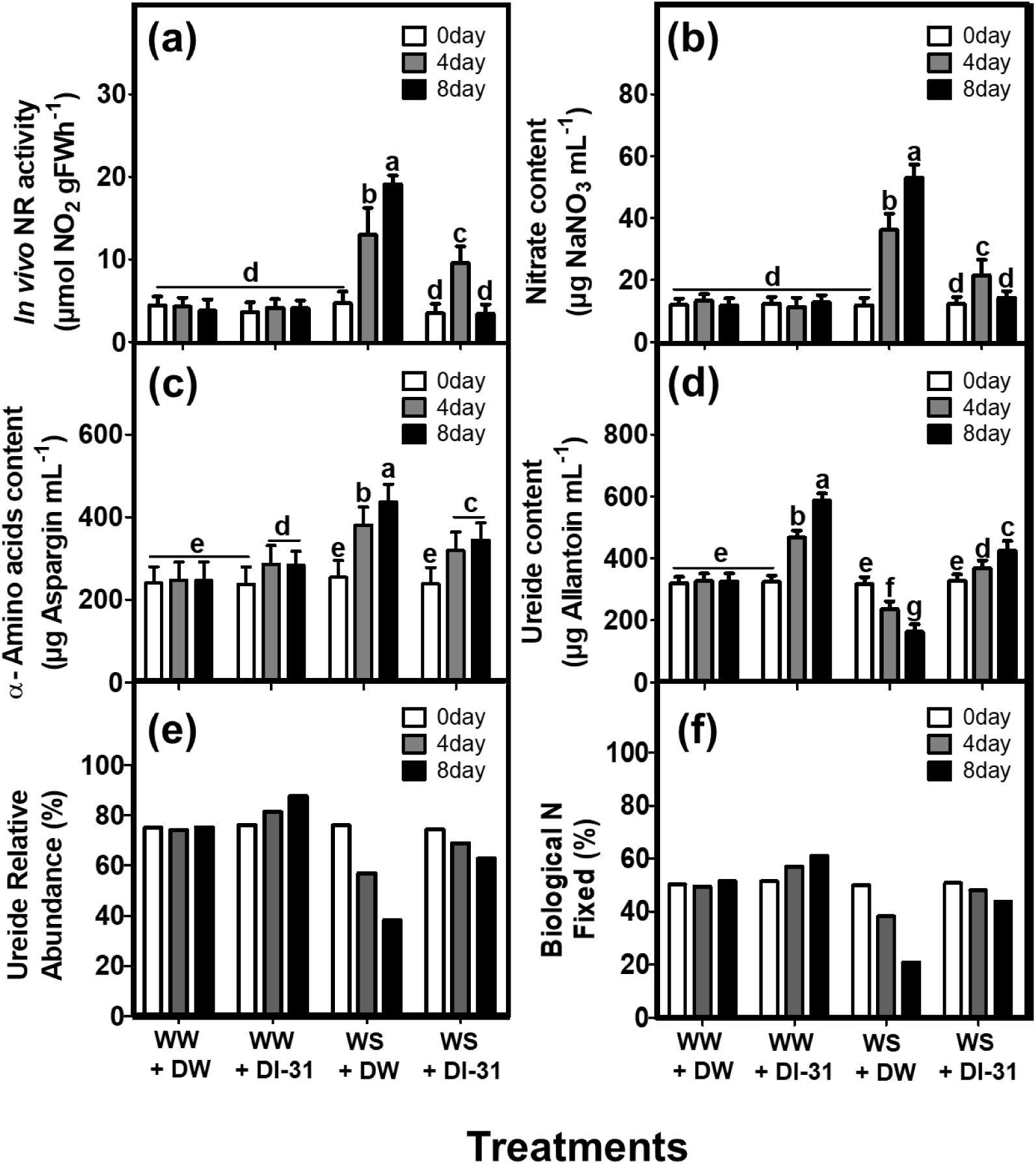
DI-31 regulates soybean nitrogen homeostasis under well-watered and water shortage conditions. Biochemical N fixation parameters such as (**a**) Nitrate Reductase (NR) *in vivo* activity, the content of (**b**) Nitrate and (**c**) α-amino acids, (**d**) ureides content and (**e**) relative abundance, and (**f**) the percentage of biological N fixed were measured in V5 soybean cv. Munasqa RR leaves. Four treatments were defined: (i) well-irrigated plants (substrate water potential (Ψs) = −0.05 MPa) sprinkled with distilled water (DW), (ii) well-irrigated plants sprinkled with DI-31, (iii) stressed plants (Ψs= −0.65 MPa) sprinkled with DW and (iv) stressed plants sprinkled with DI-31. Once the stress treatments reached the substrate water content and Ψs corresponding to moderate water stress, the DI-31 (2.23 µM) and DW treatments were performed by sprinkling to the drip point. Whole plants were collected at 0, 4 and 8 days after the application of the treatments. Data represent the mean (± SE) of two independent experiments (n= 180). Different letters on top of the bars indicate significant difference as determined by ANOVA with post hoc contrasts by Tukey’s test: (P<0.05).

## Discussion

There are two frameworks which identify drought tolerance characteristics in crop plants, including soybean (Bhatia et al. 2014). One takes into account yield variation in terms of traits affecting water use and especially the water use efficiency. The other is related to specific physiological and biochemical characteristics that lead to improvement under drought (escape, avoidance and tolerance). We took into consideration these two frameworks to discuss the applicability of DI-31 to mitigate the detrimental effects of water scarcity. As plant growth promoters, BRs participate in many developmental processes, such as cell elongation-division, assimilate translocation (Müssig 2005), the increase of shoot fresh and dry weight, plant height, petioles length and leaves size and number (Anjum et al. 2011). In this study, we corroborated that both in *A. thaliana* and soybean plants, the number of leaves, primary root length and biomass production increased due to DI-31 application; pointing out the compound capability to enhance plant photosynthetic area and growth rate. Interestingly, we observed that pot-grown soybeans treated with DI-31 and under water-limited conditions showed hydrated, expanded and young green leaves, unlike the plants subjected only to water stress that exhibited wilted, wrinkled and chlorotic phenotype. This result suggests that the compound modulates mechanism related to the dehydration postponement under water-limited conditions. Additionally, we analyzed the hydric status of Munasqa RR plants and found that well-irrigated plants, with DW and DI-31 application, evidenced a hydrated water status (91.3% average RWC). Meanwhile, the stress application reduced the RWC of DW-treated plants in a ∼28 and ∼45% at the fourth and eighth day of trial, respectively. This result corroborates the effectiveness of the stress induction by water shortage. Meanwhile, Munasqa RR plants treated with DI-31 and subjected to water shortage showed a ∼78% average RWC, and registered RWC decreases in ∼15 and ∼25% at the fourth and eighth day of stress, respectively. These results evidenced that the Munasqa RR plants subjected to stress maintained a hydrated water status due to DI-31 action.

Plant biomass is defined as the source of storable energy, mobilizable according to metabolic requirements (de Freitas Lima et al. 2017) and is highly dependent on Carbon (C) and N gathering (Zheng 2009). In a previous study, (Yamori and Shikanai 2016) correlated chlorophylls and carotenoids accumulation with an increase in dry weight production. In accordance, our results showed an increase of BMD and carotenoid levels in stressed plants treated with DI-31, strengthening the contribution of this BR analogue to energy production maintenance under water deficit conditions. There are significant correlations in soybean between the ability to cope with drought and root traits such as dry weight increase (Bhatia et al. 2014). In accordance, we observed that the DI-31 also exerts a positive effect on Munasqa RR roots, stimulating the biomass production under water shortage, which leads to an increase in water-nutrient absorption capacity to withstand the stress.

Additionally, the production of biomass represents a water cost that can be estimated through the WUE, allowing to establish a relationship between water consumption and plant production (Van Halsema and Vincent 2012). In accordance, we observed that, under water deficit, the WUE reduction was attenuated in soybean plants when DI-31 was applied. While, under conditions of proper irrigation, the application of DI-31 stimulated the WUE over time; suggesting that the compound favours the hydric status maintenance and the biomass conversion with lower water cost. Crop genetics, nutrient availability and the regulation of evapotranspiration, determine the plant WUE (Van Halsema and Vincent 2012). Besides, partial closure of stomata at a certain level of soil water deficit might lead to an increase in WUE (Miransari 2015). Plant stomata regulate key processes like CO_2_ exchange and transpiration. Thus, we examined DI-31 capacity to modulate stomatal movements, like its analogue 24-epibrassinolide, that was reported to promote stomatal closure in a dose-dependent manner (Shi et al. 2015). As we expected, DI-31 was able to induce stomatal closure in the range in which 24-epibrassinolide exerted its effect, also in a dose-dependent way. At ⁓5-10 µM concentration of DI-31, the plant stomata reached 50% of closure, in agreement with previously published results. We also demonstrated that DI-31-mediated stomatal closure is partially abolished by NADPH and peroxidase inhibitors, which suggest that this compound requires ROS production to exert its action. As part of its protective effect, DI-31-mediated stomatal closure contributes to reducing transpiration. These results are also in agreement with RWC and WUE maintenance and the hydrated phenotype observed in DI-31-treated soybean plants under water shortage. An efficient relationship between dry matter production and water consumption depends on the photosynthetic efficiency and the stomata movement that regulates transpiration (Bhatia et al. 2014). Thus, the DI-31 effect on stomatal closure, water status maintenance and roots development constituted promising indicators of the compound effect on yield potential (biomass) improvement in water-limited environments.

There are a plethora of physiological and biochemical traits involved in the water stress response. In this paper, we decided to evaluate the effect of DI-31 on stress response markers associated with osmotic adjustment, antioxidant activity, respiratory burst and chlorophyll and photoprotective pigments accumulation. BRs can stimulate ROS production, which acts as second messengers in several processes like photosynthesis, respiration and tolerance/resistance to environmental stresses (Tripathy and Oelmüller 2012). Cells tightly control ROS level to prevent oxidative injuries, partially through enzymatic antioxidants like SOD and a wide range of peroxidases among which stand out APX and POX enzymes (Sharma et al. 2012). We observed that DI-31 foliar application triggers *A. thaliana* respiratory burst. Besides, under well-watered conditions the compound caused an increased in SOD, APX and POX activities, in *A. thaliana* and soybean cv. Munasqa RR plants. While, under water deficit, DI-31 effect on antioxidant enzymes was even more pronounced. These findings indicate the compound intrinsic ability to activate mechanisms like oxidative burst, ascorbate-glutathione (Asa-GSH) cycle and phenols synthesis. We analysed lipid peroxidation through the MDA level and found that water stress, as expected, provoked an increase in MDA content. However, the MDA formation decreased in stressed plants due to DI-31 treatment, suggesting that the compound diminish the lipid-peroxides accumulation. Related to osmotic adjustment, we quantified an increase of free proline content in Munasqa RR plants subjected to DI-31 treatment, especially under water shortage conditions. In response to water deficit, plants can activate or increase a major defence mechanism composed, among others, by enzymatic antioxidants (Sajedi et al. 2011). Thus, a high antioxidant capacity is linked to increased crop stress tolerance (Sharma et al. 2012). Overall, our data indicate that the DI-31 promote the degradation of O_2_^−^, H_2_O_2_ and lipoperoxidation products under stress. Additionally, in soybean plants under water scarcity, the compound also contributes to chlorophyll content maintenance and carotenoids and free proline overproduction. These findings agree with the obtained in wheat, rice and potato cultivars, where the accumulation of non-enzymatic antioxidants and the lipid-peroxides reduction resulted in a higher survival rate, yield and tolerance to drought/osmotic stresses (Amini et al. 2015).

BRs pleiotropic effects in plant growth and development, as well as resistance/tolerance to biotic and abiotic stresses, offer exciting potentialities for enhancing crops productivity and quality (Ali and Ashraf 2008; Baghel et al. 2019; Gill et al. 2017). Nevertheless, few studies address the effect of these hormones on the balance of macronutrients such as N, even though (Shu et al. 2016) reported that foliar application of 24-epibrassinolide significantly enhanced the activity of assimilative N2-fixation critical enzymes in tomato plants. While, (Wang et al. 2019) suggest that the transcriptional factor BZR1, a BRs positive regulator, possibly plays a critical role in tomato N-starvation response.

It’s known that plant N regulation is needed to maintain optimum photosynthetic rate and therefore biomass production since most of leaf N is a constituent element of the C assimilation protein ribulose 1,5-bisphosphate carboxylase/oxygenase (RuBisCO) strongly involved in photosynthesis (Rotundo and Cipriotti 2017; Sinclair and Horie 1989). Soybean demands high N concentrations, and it is often grown on soils with low N availability thus the biological N2 fixation (BNF) makes significant contributions to the plant growth, yield and high-protein seed and forage production (Peoples et al. 2009). Few reports exist of low BNF contributions in soybean, and most of these results come from breeding and cropping-managements with high inputs of N-fertilizers (Hungria and Mendes 2015).

Soybean BNF depends on the formation of the nodule (Denton et al. 2017). Interestingly, soybean nodules are determined; there is no permanent meristem, so its growth depends on expansion instead of cell division. In consequence, the nodular N-fixing capacity depends on the nodules size and the number of bacteroids (fixing centres) located in the infected medulla (de Felipe Antón 2007). The nodule development and therefore the BNF are strongly related and also are severely affected during periods of moisture deficiency (Purcell et al. 2004; Sinclair et al. 2007); mainly through ROS and/or reactive N species (RNS) accumulation leading to the Leghemoglobin self-oxidation and the Nitrogenase complex inactivation (Puppo et al. 2005). Water deficit also provokes nodule dehydration and wrinkling, with a marked reduction of the infected central medulla and finally the nodular senescence activation (Hernández-Jiménez et al. 2002). Our findings demonstrate that the DI-31 retarded the active nodules senescence. The presence of numerous light pink nodules, after eight days of water scarcity, allowed us to assume that the DI-31 reduced the Leghemoglobin self-oxidation; either by activating (i) ROS/RNS-scavenging or exclusion mechanism, (ii) water economy or (iii) Leghemoglobin recycling pathways. On the other hand, the shrinking in the cortex area, observed in all the DI-31-treated nodules, might be explained by the expansion of the infected central medulla, which suggests the compound ability to increase the bacterial colonization area and therefore the nodules N-fixing capacity. Moreover, the significant periderm thickness, quantified in DI-31 treated nodules under stress, indicates an outer layers reinforcement to cope with water deficit.

To complement the results obtained in the nodulation morphology analysis, we evaluated several N homeostasis biochemical markers and found that DI-31 regulates the N homeostasis mechanism, especially under stress. According to (Rodríguez-Navarro et al. 2011), the Nitrate Reductase (NR) is the first enzyme in nitrates assimilative reduction (NAR) pathway, catalyzing the nitrate (NO3^-^) conversion into nitrite (NO_2_^-^) which is subsequently transformed in ammonia (NH_3_) and then in assimilable ammonium (NH_4_^+^). The NR is synthesized and degraded continuously, so the control of its activity is through a substrate regulation (Rajasekhar and Oelmüller 1987). Thus high levels of NO_3_^-^ increase the enzyme activity. Our results show that soybean plants under water scarcity present high levels of nitrate and NR activity. Therefore, plants NAR was possibly stimulated to satisfy the N demand caused by the nodular senescence and BNF deficit due to stress occurrence. On the contrary, stressed and DI-31 treated plants showed a reduced loss of active nodules, lower NR activity and nitrate content, so we speculate that the DI-31 also modulates the mineral N absorption pathways.

In soybean, the NH_4_^+^ resulting from the symbiosis is converted into ureides, which are synthesized in the nodules and transported to the leaves through the xylem (King and Purcell 2005; White et al. 2007). In contrast, the NH_4_^+^ produced by NAR is converted into α-amino acids, mainly asparagine (White et al. 2007). The α-amino acids accumulation observed in Munasqa RR leaves, due to water shortage, agrees with an increase in NAR pathway indicators. On the other hand, ureides have four C and four N atoms in their chemical structure, therefore are more efficient for N transport than α-amino acids, which only have two N atoms. Besides, the nodules require high amounts of C, before an active BNF, which agrees with the ureides efficiency in transporting N atoms equivalent to their C number (Freixas et al. 2010). Nitrogen metabolism under water deficit plays a crucial role in the BNF regulation, which occurs through a negative feedback mechanism (King and Purcell 2005). It is not clear if the accumulation of ureides in plants leaves constitutes a stress tolerance or susceptibility response. (Vadez and Sinclair 2002) report a decrease in Nitrogenase activity and the content of ureides in the leaves in drought-sensitive soybeans subjected to water scarcity and manganese treatments.

Meanwhile, a research performed by (King and Purcell 2005) also in drought-susceptible soybeans correlates a BNF decrease with the ureides accumulation in leaves. In a subsequent study, the same authors state that there is no evidence of tolerance when soybean varieties, sensitive or tolerant to water deficit, increase the ureides concentration in roots and decrease in leaves during drought (King and Purcell 2005). Ureides accumulation in the nodule directly inhibits the BNF during stress (Charlson et al. 2009), therefore, if their proper transport from the nodules to leaves is guaranteed, the BNF inhibition could be prevented, at least temporarily. Thus, the increase in ureide content and relative abundance in Munasqa RR leaves due to DI-31 application, could contribute to the BNF maintenance under water scarcity. (Santachiara et al. 2018) reported that BNF in soybean represents ⁓60% of the total N uptake. Whereas, in Argentina, the range of N derived from BNF varied from 46 to 71% in farmers’ fields (Collino et al. 2015) . In agreement, our results demonstrate that the percentage of N fixed biologically in Munasqa RR plants, covers ⁓50.4% of the total N demand. Interestingly, the significant BNF increase in well-watered and stressed plants, both treated with DI-31, reinforced the hypothesis of probable BRs-plant-nodule crosstalk that positively modulates the N homeostasis.

Authors like (Sinclair and Horie 1989), (Rotundo and Cipriotti 2017) and (Santachiara et al. 2018) actively discuss that N plays a central role in the proteins homeostasis, photosynthesis, respiration, leaf area generation and water use efficiency. Thus, we considered that the DI-31 action on NAR and BNF could link with other physiological traits such as photosynthesis, antioxidants activation, water preservation, stomatal movement, growth and biomass production/duration, previously discussed in the paper. The source-to-sink N partitioning directly influences the grain production, and soybean shows a strong positive correlation between seed yield and N uptake (Rotundo and Cipriotti 2017; Salvagiotti et al. 2008; Tamagno et al. 2017). In accordance, it is conceivable to speculate that the DI-31 application, beyond the short and middle-term effect on the growth, respiration, anti-stress response and N homeostasis, might have also a long-term effect on soybean seed quality and yield.

## Conclusions

The exogenous application of DI-31 stimulates leaves and roots development, photosynthetic and water/nutrient absorption capacity, water economy partially through stomatal closure induction, WUE and biomass production and duration. Also, the compound promotes respiratory burst, osmotic adjustment, the synthesis of photoprotective pigments and enzymatic antioxidants, improving the ROS-scavenging and preventing the accumulation of damaging products like MDA. Moreover, the compound showed a remarkable protective effect in soybean nodulation and N homeostasis. Accordingly, plants treated with DI-31 showed a higher number of active nodules, with a larger size, reinforced periderm and larger medulla, as well as higher BNF values under stress. Thus, we propose that DI-31 represents a practical value as a promising bio-stimulant that might help to alleviate stress-derived impacts on soybean production. Moreover, the potential use of DI-31 to promote growth and regulate stress-response could represent a sustainable and environmentally safe alternative for integrative crop resilience management amidst climate change threats.

## Acknowledgements

This work was supported by a grant from the National Council of Scientific and Technical Research (CONICET) of the Argentine Republic (Res.3224 to LSPB), a Bioplantas Institutional Project (0018 to LSPB) and GrB2/GrB3 EEAOC Grains Program work plans (to EMP). The authors would like to especially thank Dr Christian Wilhelm Bodo Bachem from the Department of Plant Sciences of Wageningen University and Research, for his very accurate and creative corrections and suggestions.

